# Oriented evening flight behaviour in the Bogong moth revealed through automated video tracking

**DOI:** 10.1101/2022.05.19.492638

**Authors:** Jesse R A Wallace, David Dreyer, Jochen Zeil, Eric J Warrant

## Abstract

During their period of summer dormancy, Australian Bogong moths *Agrotis infusa* undertake seemingly random evening flights, filling the air with densities in the dozens per cubic metre. The purpose of these flights is unknown, but they may serve an important role in Bogong moth navigation, which remarkably enables them to return to the same exact summer sites— generation after generation—after migrating around 1000 km, and with no opportunity to learn their route or destination from prior generations. The recent development of the camera-based insect monitoring method, Camfi, enables quantitative observations of Bogong moth behaviour at an unprecedented scale. To gain a better understanding of the summer evening flights of Bogong moths, we have extended Camfi to facilitate automated video tracking of flying insects, taking the already-high throughput of the method to a new level. We used this new method to record the evening flight behaviour of Bogong moths in two elevational transects below the summit of Mt. Kosciuszko, NSW, on a single night in February 2021, and found that these flights were not random, but were systematically oriented in directions relative to the azimuth of the summit of the mountain. These results stimulate interesting and plausible hypotheses relating to previously unexplained summer evening flight behaviour of Bogong moths, and the mechanisms of their long-distance navigation.

## 1 Introduction

During their period of summer dormancy, known as aestivation, Australian Bogong moths *Agrotis infusa* remain huddled within cool, dark crevices of granite outcrops that dot the peaks of the Australian Alps, tiling the walls with an estimated density of up to 17,000 moths per square metre (Common, 1954). However, during evening twilight, the moths are known to emerge from their hiding spots and undertake seemingly random flights (Common, 1954; Wallace et al., 2021; Warrant et al., 2016). Although these flights are only undertaken by a portion of the moths at a particular site, they are enough to fill the air with densities probably reaching dozens per cubic metre (Wallace, personal observations).^1^ The purpose of these flights is unknown, although observations have been made of Bogong moths using them to visit water to drink (Common, 1954; Warrant et al., 2016).

It could be that the evening flights of aestivating Bogong moths are used as a sort of learning flight to calibrate their navigational machinery, akin to how homing insects familiarise themselves with an area of interest (Collett and Zeil, 2018), or how night-migratory birds calibrate their star compasses and other compass systems prior to migration (reviewed by Foster et al., 2018; Pakhomov and Chernetsov, 2020). Alternatively, one might hypothesise that these flights are undertaken by moths who are dissatisfied with their resting place—perhaps being too warm or not dark enough— and are seeking a more favourable site in which to continue their aestivation. The former possibility may be necessary, particularly as the departure date for the return migration draws near. We might expect the latter possibility to play a more important role during the first few months of aestivation, as Bogong moths are known to occupy higher and higher elevation sites as the summer progresses (Green, 2003). Disentangling these possibilities is important for our understanding of the mechanisms of Bogong moth navigation, which remarkably enable them to return to the exact same summer sites—generation after generation—following a migration of around 1000 km, having had no opportunity to learn their route or destination from prior generations.

The recent development of Camfi, a camera-based system for monitoring evening flight behaviour in wild Bogong moths (Wallace et al., 2021), presents an opportunity to make quantitative observations of the dynamics of the behaviour with an unprecedented scale and spatiotemporal resolution. In this paper, we demonstrate how observations made using Camfi can begin to disentangle the causes of Bogong moth evening flights, and we provide evidence of directed flights undertaken by the moths near aestivation sites, giving clues as to the purpose of the flights.

In order to measure the direction of flight, we have extended Camfi to facilitate automated video tracking of insects flying above the camera. Full details of this new method are presented in this paper, and the method has been implemented as part of the Camfi package, freely available from https://github.com/J-Wall/camfi.

## 2 Methods

### 2.1 Detection of flying Bogong moths using Camfi

Wallace et al. (2021) introduced Camfi, a method for monitoring the activity of flying insects using still images obtained from off-the-shelf wildlife cameras. Camfi has been used to monitor the activity of migratory Bogong moths arriving at, aestivating in, and departing from their summer range in the Australian Alps (Wallace et al., 2022). Camfi performs automatic detection of flying insects using the Mask R-CNN framework (He et al., 2017), and at the time of writing, has been trained on a set of 4901 manually annotated images of flying Bogong moths (Wallace et al., 2022). In addition, Camfi automatically measures wingbeat frequency of detected insects in still images, which is useful for assigning species identity to observations of flying insects.

An advantage of Camfi is its flexibility with regard to the temporal resolution of data collection. Depending on the research question, cameras can be set to capture an image at relatively long intervals, on the order of minutes, or they can be set to capture images at a very high rate, which in the case of video clips is on the order of hundredths of a second (typically 25-30 frames per second). However, when analysing Camfi data which have been obtained from high-rate captures (namely, videos), individuals will be detected multiple times, since each moth will be seen in each of many consecutive video frames as they pass by the camera. This results in detection counts being inflated by insects which have lower angular velocities relative to others from the perspective of the camera, and therefore spend more time in-frame. Therefore, to facilitate the use of videos by Camfi, we need to be able to track observations of individuals in a sequence of video frames, so we can count each individual only once.

In the following sections, we introduce an extension to Camfi which enables analysis of video data. This includes proper handling of video files, as well as tracking of individuals through consecutive frames. In addition to ensuring individual insects are only counted once per traversal of the camera’s field of view, the new method allows for measurement of the direction of displacement of insects as they travel through the air.

### 2.2 Multiple object tracking

Multiple object tracking is a challenging problem which arises in many computer vision applications, and which has been approached in a variety of different ways (reviewed by Luo et al., 2021).

A common approach to multiple object tracking is “detection-based tracking” (also known as “tracking-by-detection”), in which objects are detected in each frame independently, and then linked together using one of a number of possible algorithms. Typically, this requires the use of a model of the motion of the objects to be tracked, along with a method which uses the model of motion to optimise the assignment of detections to new or existing trajectories. In many approaches, the modelled motion of tracked objects is inferred by combining information about the position of the objects in multiple frames. An obvious challenge arises here because the model of motion requires reliable identity information of objects detected in multiple frames, whereas the identity of the objects usually must be inferred from their motion (including their position). This circular dependency—between the inference of object identity and the model of motion—can be dealt with in a number of ways, including probabilistic inference via a Kalman filter (Reid, 1979) or a particle filter (e.g. Breitenstein et al., 2009), or through deterministic optimisation using a variety of graph-based methods.

Our approach to multiple object detection removes the requirement of an explicit model of motion entirely, by utilising two peculiar properties of the Camfi object detector. The first of these properties is that the Camfi detector obtains information about the motion of the flying insects it detects from the motion blurs the insects generate, which it stores in the form of a polyline annotation.^2^ Since this information is obtained from a single image, and therefore a single detection, it does not depend on the identity of the insect, solving the previously mentioned circular dependency problem. The second property is that the Camfi detector is robust to varying exposure times, owing to the fact it has been trained on images with a variety of exposure times. This in turn means that the detector is robust to the length of the insects’ motion blurs. Ultimately, these two properties, along with the fact that the insects appear as light objects on a dark background, mean that it is possible to use the Camfi detector to make a single detection of an individual insect traversing multiple consecutive frames. Thus, the trajectories can simply be formed using bipartite graph matching of overlapping polyline annotations, using only information provided by the detections themselves, using the method described in section 2.3.

### 2.3 Automated flying insect tracking

The algorithm described in this section has been implemented in Python, and is packaged together with the Camfi software, available under the MIT licence from https://github.com/J-Wall/camfi.

We will now describe our algorithm for tracking flying insects in short video clips. The algorithm uses a detection-based tracking paradigm, relying heavily on the Camfi flying insect detector described by Wallace et al. (2021). Accordingly, we will not present a detailed explanation of the detector here, but will instead focus on the process of linking detections into trajectories. Readers interested in the details of the detector should consult Wallace et al. (2021).

An example of the sequence of steps taken by the tracking algorithm described in this section is illustrated in Fig. 1. For brevity, the example shows the algorithm operating on three frames only, however the algorithm can operate on any number of frames, up to the memory constraints of the computer it is running on.

**Figure 1:**
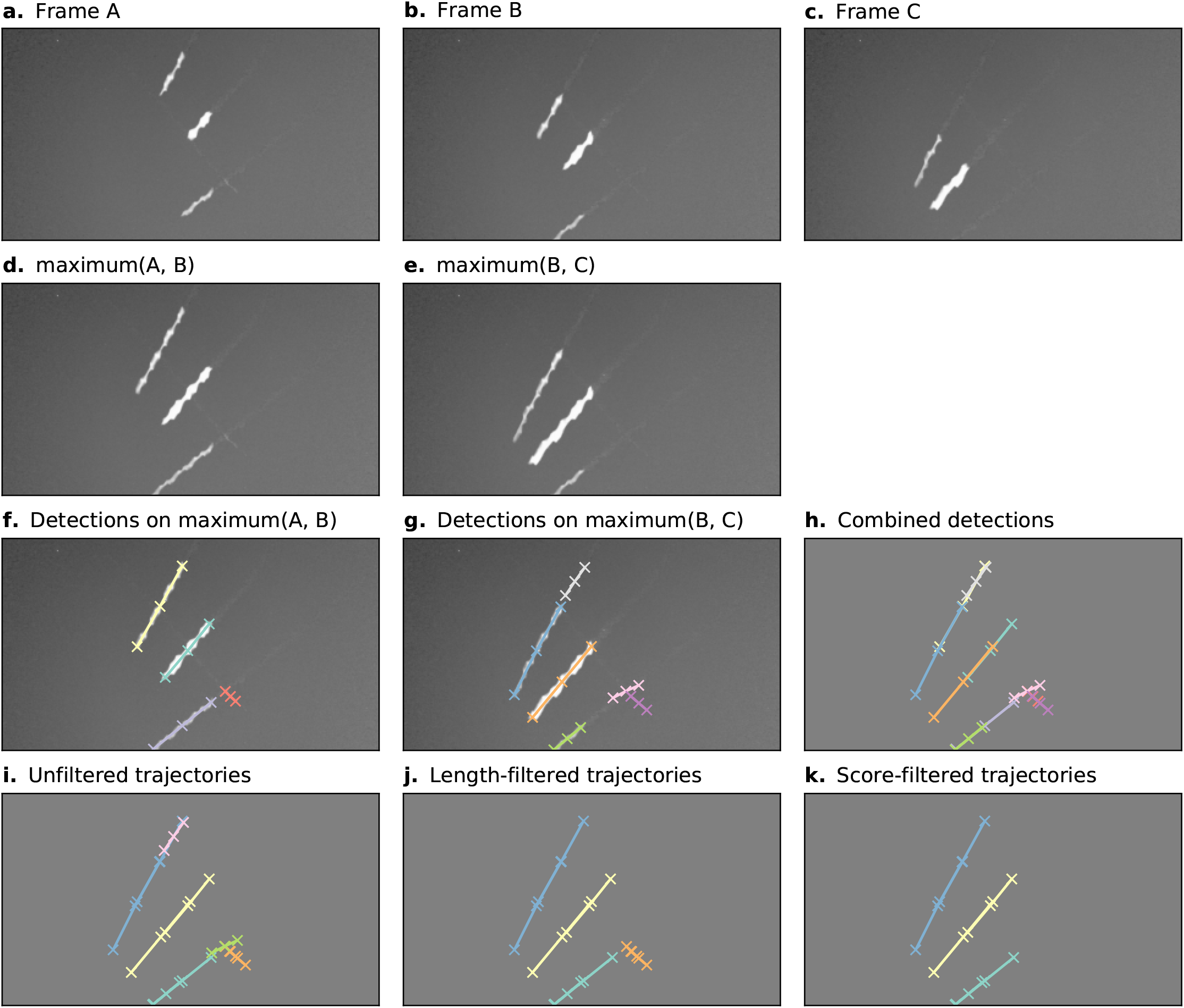
Automatic annotation is performed by Camfi on the maximum image of each pair of consecutive frames, allowing trajectories to be built from overlapping detections. Here, an example of this process is shown for three consecutive video frames. **a–c**. Three consecutive video frames containing multiple flying insects. **d–e**. The maximum image of each sequential pair of frames. f–g. Flying insects are detected in the two-frame maximum images using Camfi. h. Detections from f and g together on a plain background. i. Detections from sequential time-steps are combined into trajectories using bipartite graph matching on the degree of overlap between the detections. **j**. Trajectories containing fewer than three detections are removed. **k**. Finally, trajectories are filtered by mean detection score (trajectories with mean detection score lower than 0.8 are removed).

First, the video frames are prepared for flying insect detection. A batch of frames is loaded into memory (e.g. Fig. 1a–c, although typically this would be a video clip). The maximum image of each sequential pair of frames is then calculated by taking the maximum (brightest) value for each pixel between the two frames (Fig. 1d–e). This produces images with lengthened motion blurs of the in-frame flying insects, approximating the images which would be obtained if the exposure time of the camera were doubled. Importantly, the motion blurs of an individual insect in consecutive time-steps overlap each other in these maximum images.

Detection of flying insects is performed on the maximum images using the Camfi detector (Wallace et al., 2021), producing candidate annotations of insect motion blurs to be included in trajectories (Fig. 1f–g). The Camfi detector produces polyline annotations which follow the respective paths of the motion blurs of flying insects captured by the camera. Because the motion blurs of individual insects overlap in consecutive frames, so too do the annotations of those blurs (e.g. Fig. 1h). This enables the construction of trajectories by linking overlapping sequential detections.

Detections in successive time-steps are linked by solving the linear sum assignment problem using the modified Jonker-Volgenant algorithm with no initialisation, as described by Crouse (2016). In order to do this, a formal definition of the cost of linking detections is required. We call this cost the “matching distance,” which we denote by *d*_*M*_. Consider two polyline annotations *P*_*a*_ and *P*_*b*_, which are sequences of line segments defined by the sequences of vertices 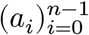 and 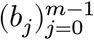 respectively, where *a*_*i*_, *b*_*j*_ *∈ ℝ*^2^. We define *d*_*M*_(*P*_*a*_, *P*_*b*_) as the second smallest element in {*d*(*a*_0_, *P*_*b*_), *d*(*a*_*n*−1_, *P*_*b*_), *d*(*b*_0_, *P*_*a*_), *d*(*b*_*m*−1_, *P*_*a*_)}, where *d*(*x, P*) is the Euclidean distance from a point *x ∈ ℝ*^2^ to the closest point in a polyline *P* ⊂ *ℝ*^2^. This definition of *d*_*M*_ is efficient to compute, and allows us to discriminate between pairs of detections which come close to each other by chance (perhaps at very different angles) and pairs of detections which closely follow the same trajectory (i.e. roughly overlap each other).

After solving the assignment problem, a heuristic is applied to reduce spurious linking of detections into trajectories, where links with *d*_*M*_ values above a specified threshold are removed. Trajectories are built across the entire batch of frames by iteratively applying the detection linking procedure for each consecutive pair of time-steps (Fig. 1i). Trajectories containing fewer than three detections are removed (Fig. 1j), as are trajectories with low mean detection scores (Fig. 1k). We used a mean score threshold of 0.8 to produce the final set of trajectories for further analyses. When analyses relating to flight track directions are required, we apply an additional filtering step to constrain analysis to detections inside a circular region of interest within the frame. This eliminates directional bias arising from the non-rotationally-symmetrical rectangular shape of the video frames.

Diagnostic plots of tracking performance over an entire short video clip can be made by taking the maximum image of the entire video clip, and plotting the detected trajectories as a single image using a different colour for each trajectory (e.g. Fig. 2). For example, we can see good performance of the tracking procedure in Fig. 2a, where all trajectories except one appear to have been correctly built. The one exception is an insect close to the centre of that figure which appears to have had its trajectory split in three parts (seen as three different coloured segments), most likely due to occlusion by another insect. Fig. 2b shows the result of constraining these trajectories to a circular region of interest to remove directional bias (in this case, this happened to solve the aforementioned split trajectory, but only by coincidence—the orange and purple tracks were removed for overlapping the edge of the circle).

**Figure 2:**
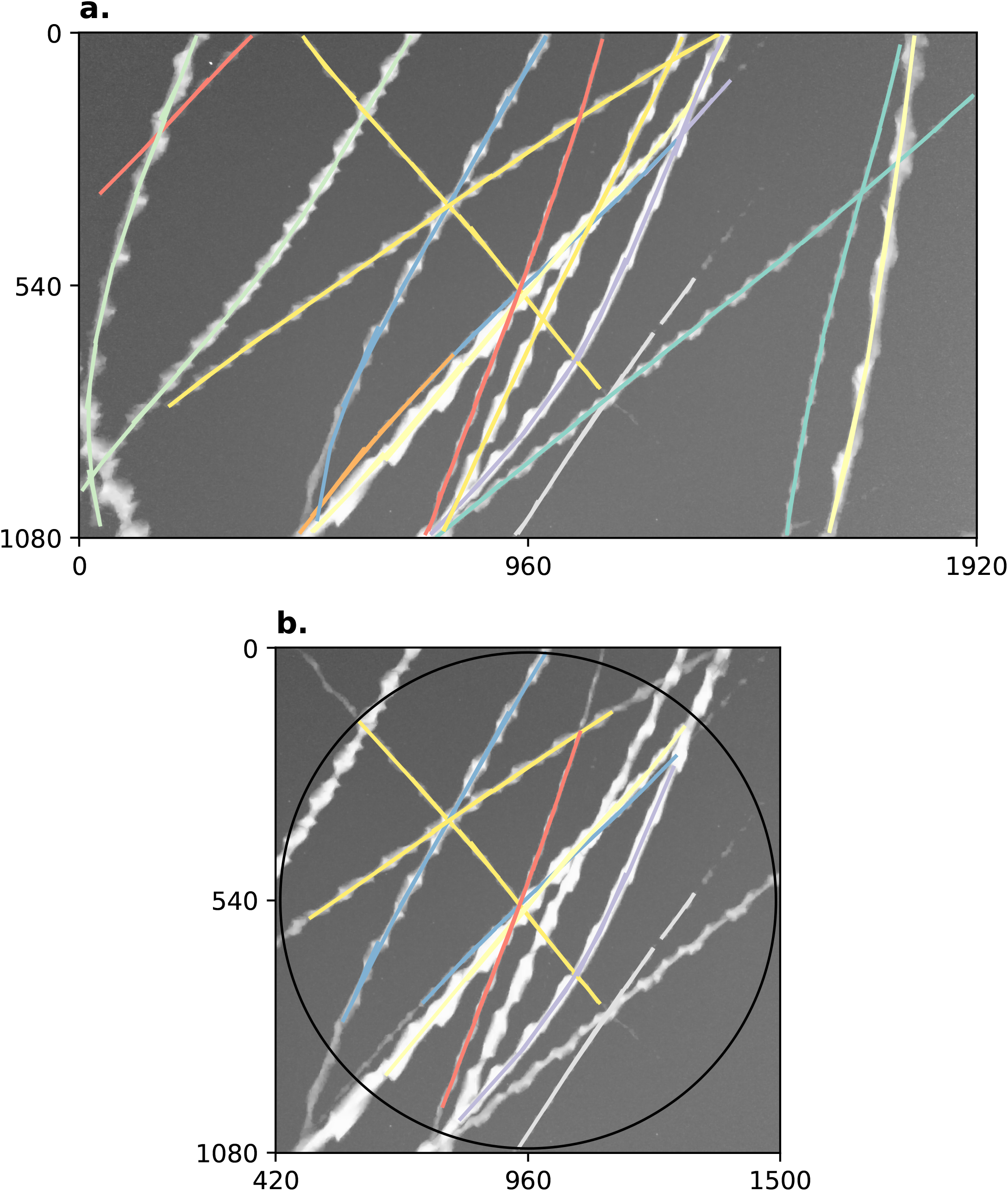
Example summary of trajectories followed by insects flying past a camera during a 5 s video clip. Axes on both plots show pixel row and column numbers. **a**. Maximum (brightest) value of each pixel across every frame in the clip with annotations overlaid. Visible bright streaks are made by the motion blurs of Bogong moths flying past the camera. The colour of an annotation indicates its membership in a unique trajectory, as predicted by our method. **b**. Annotations constrained to circular region of interest. Using only these trajectories eliminates directional bias resulting from the non-rotationally symmetrical rectangular shape of the frame. Black circle shows region of interest.

### 2.4 Camera placement and settings

A total of ten cameras (BlazeVideo, model SL112) were placed in two transects below the summit of Mt Kosciuszko, NSW on the afternoon of 18^th^ February 2021, and collected the following morning. The first transect, which we call `kosci_south`, was placed on the south-eastern slope, running from the shore of Lake Cootapatamba up to the Kosciuszko South Ridge, and ranging in elevation from 2046 m to 2151 m. The second transect, which we call `kosci_north`, was placed on the north-western slope below the summit, with five cameras ranging in elevation from 2050 m to 2220 m. The positions of each camera are shown in Fig. 3a. The `kosci_north`1 and `kosci_north`2 locations were both within 10 m of known Bogong moth aestivation sites.

**Figure 3:**
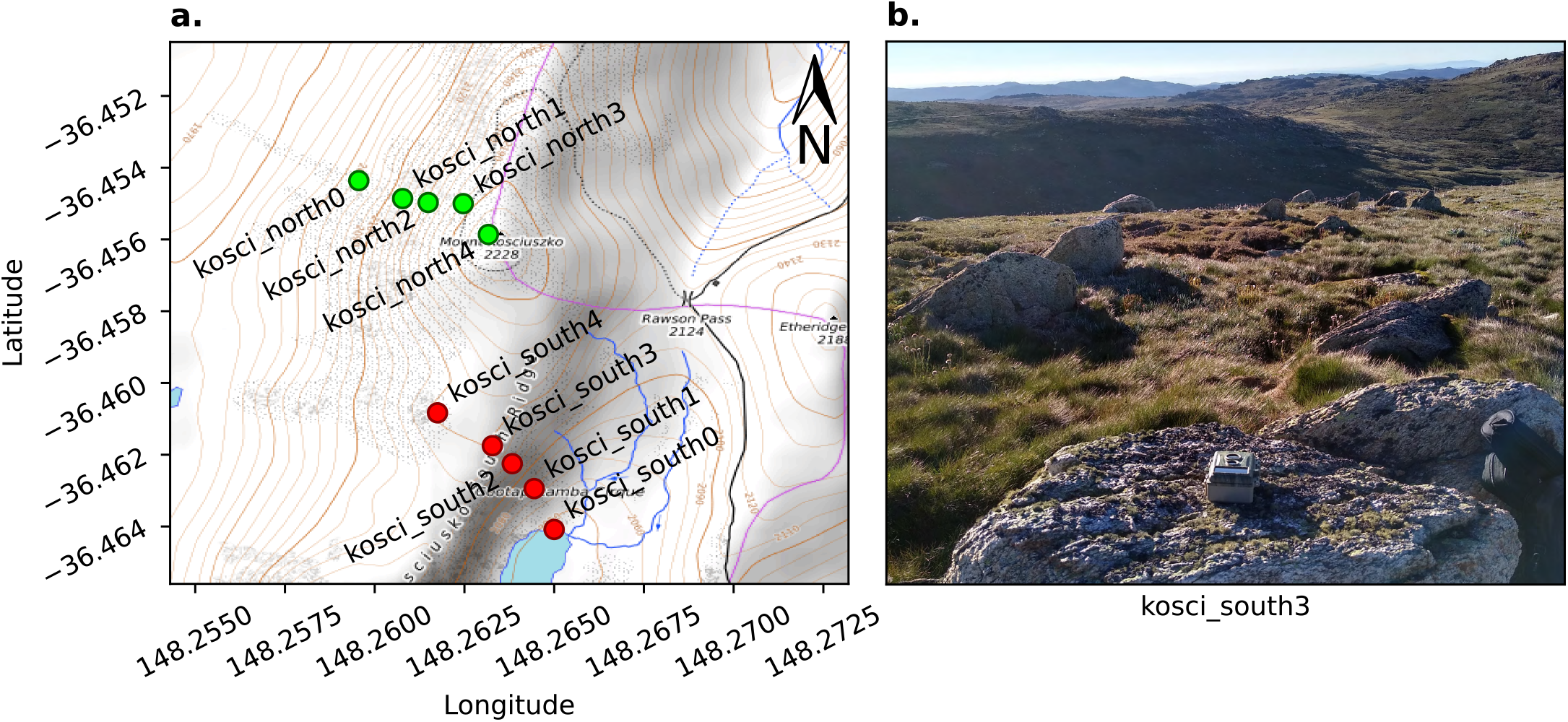
Cameras were placed on two transects on the slopes of Mt Kosciuszko, NSW. **a**. Map of camera locations. Contour lines show 10 m changes in elevation. The transects were `kosci_south` (*red*), on the south-eastern slope towards Lake Cootapatamba and `kosci_north` (*green*), on the north-western slope, below the summit. **b**. Example placement of camera at `kosci_south`3 location. Map data available under the Open Database Licence at openstreetmap.org. © OpenStreetMap contributors, SRTM. Map tiles credit: © OpenTopoMap (CC-BY-SA).

The cameras were placed such that their lenses pointed up into the sky, and the compass orientation of each camera was noted so that analysis of flight direction could be performed. Fig. 3b shows an example of the placement of one of the cameras. The cameras were set to take an image, along with a 5 s video clip every 30 s for the duration of the evening. Illuminance of the clear sky was recorded at multiple time points during evening twilight, near the `kosci_north`2 location (Fig. 3a) using a digital luxmeter (Hagner, model E4-X). Luminance was also recorded from the rock face and a white standard in a few locations inside and outside a Bogong moth aestivation cave, also near the `kosci_north`2 location, using a digital photometer (Hagner, model ERP-105).

### 2.5 Computational analyses

Flying insects were detected in the 5 s video clips using Camfi (Wallace et al., 2021) and tracked using the method described above (section 2.3). Track directions were modelled with the orientation models described by Schnute and Groot (1992) using the CircMLE R package (Fitak and Johnsen, 2017). Maximum likelihood models were selected using Akaike’s information criterion (AIC, Akaike, 1973).

## 3 Results and Discussion

At approximately 20:30 Australian Eastern Daylight Time (AEDT; UTC+11:00) the cameras switched to night mode and started using their infra-red flash. Video clips taken before this time were omitted from analysis as it was found that detection was unreliable for video clips taken in day mode. A total of 6,515 night-mode video clips were recorded, and from these 11,147 flying insects were detected. The vast majority of these are likely to be Bogong moths, as we observed a large number of them (and no other species) flying close to our vantage point near `kosci_north`2 throughout the evening. Sky illuminance varied from 106.5 lx to 0.0132 lx over the course of evening twilight (Fig. 4b, *red trace*).

**Figure 4:**
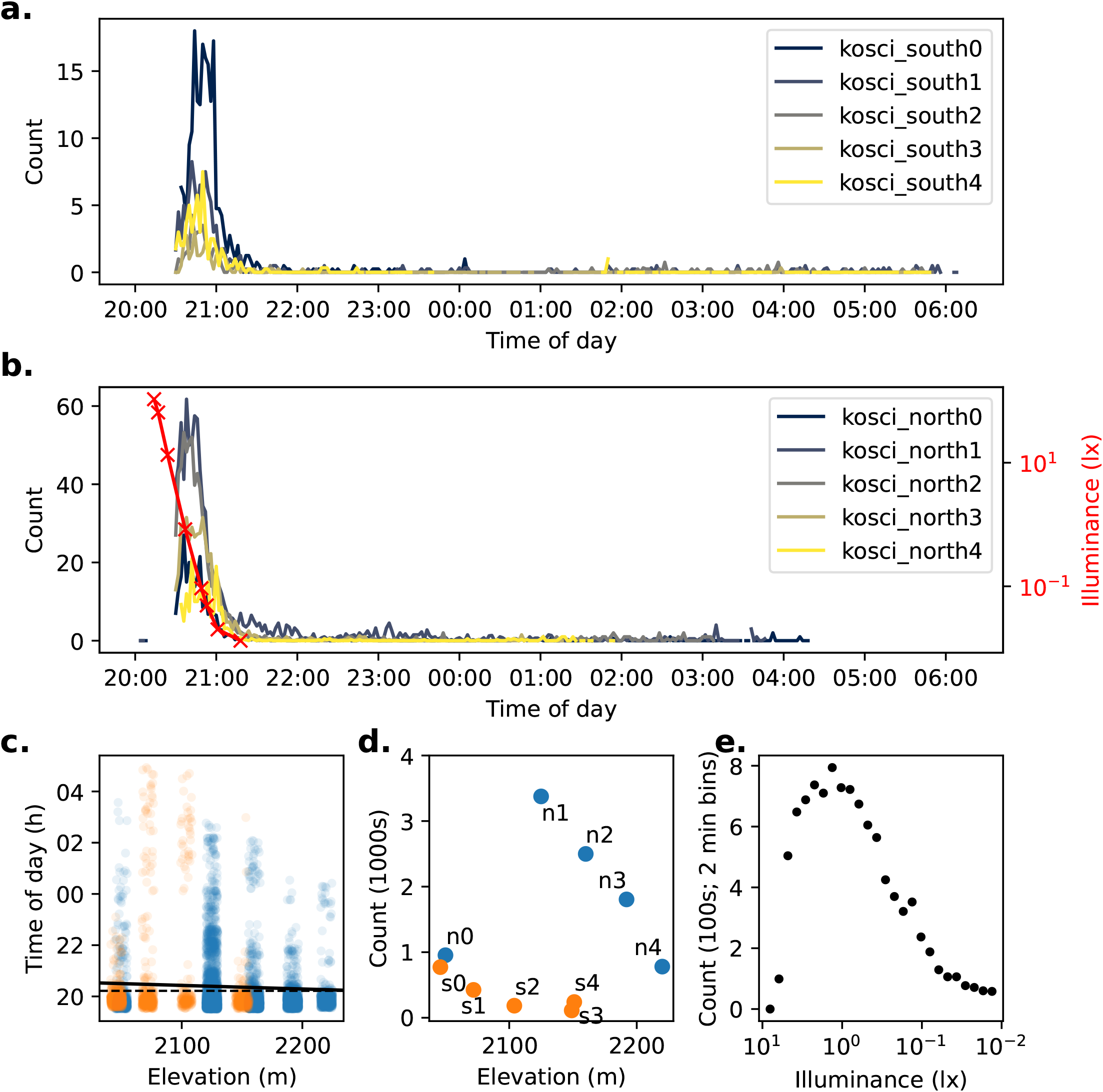
Summary of detections of flying insects on two transects of Mt. Kosciuszko on 18^th^–19^th^ February 2021. **a–b**. Time series of counts of flying insect trajectories detected at each location in `kosci_south` (*a*) and `kosci_north` (*b*) transects. Counts have been smoothed by taking the average over 2 min bins. Illuminance readings recorded from the sky close to the `kosci_north`2 site are shown in *b* (*red trace*). **c**. Time of each detection by elevation. Jitter (random fluctuation) is applied to elevation values to assist readability. A linear regression of time against elevation is shown by *solid black line* (*R*^2^ = 0.0065, slope = −5.59 s/m). Despite the small effect size, the slope is statistically significant (Wald test, *p* = 1.8 × 10^−17^; null “zero slope” hypothesis is indicated with *dashed black line*). **d**. Total number of detections at each location by elevation. Location names are labelled “n0” = “`kosci_north`0,” “s3” = “`kosci_south`3,” etc. Points in *c* and *d* are marked according to transect; *blue* = `kosci_north`, and *orange* = `kosci_south`. **e**. Total detection count in 2 minute bins (pooled across all locations) plotted against illuminance (log-linearly interpolated from recorded measurements; *b, red trace*). Measurements taken between 20:28 and 21:18 are included.

### 3.1 Activity levels

A strong peak in activity was observed during evening twilight at all sites on both transects. Activity plummeted just before 21:00 AEDT, coinciding with the end of nautical twilight (Fig. 4a–b). Outdoor illuminance dropped from about 1 lx during the activity peak to below 0.1 lx after activity had plummeted (Fig. 4e). Of the 11,147 total flying insect detections, 8,589 occurred before 21:00 AEDT (from 576 video clips), and 10,163 occurred before 21:30 AEDT (from 1,176 video clips; coinciding with the end of astronomical twilight). This agrees with previous observations of Bogong moth flight activity exhibiting large peaks during evening twilight (Common, 1954; Wallace et al., 2022, 2021; Warrant et al., 2016). The activity peak was most pronounced at the `kosci_north`1 and `kosci_north`2 locations (Fig. 4d), presumably owing to the proximity of these locations to known Bogong moth aestivation sites.

Previous work has demonstrated that occupied Bogong moth aestivation sites vary in elevation over the course of the summer, with moths occupying higher elevation sites as summer progresses and temperatures increase (Green, 2003; Wallace et al., 2022). There are also some indications that towards the end of summer—as temperatures start to drop—Bogong moths may re-occupy lower altitude sites, possibly as they start their return migration to the north, north-east, and east (Common, 1954). Clearly, these patterns of site occupation require movement of individuals between elevations. Since the mode of Bogong moth locomotion is predominantly flight, and they are known to be particularly strong flyers (Dreyer et al., 2018; Warrant et al., 2016), we would expect these movements between elevations to occur over short time periods—on the order of minutes or hours, rather than the days or weeks that previous monitoring methods have measured (e.g. direct observation (Caley and Welvaert, 2018; Common, 1954); fox scats (Green, 2003); daily-pooled still-image camera monitoring (Wallace et al., 2022)).

If there was a trend for Bogong moths to move between elevations on the night of our recordings, then we would expect the timing of the peak in detections to vary with elevation. However, this type of temporal shift may be difficult to detect, especially if the duration of travel for an individual moth between the elevations is short with respect to the total duration of the detection peak (since, in that case, most of the variation in detection time would be explained by variation in the times that moths take flight, rather than movement of moths across an altitudinal gradient). Fortunately, the present method provides tremendous statistical power to detect such weak interactions, owing to the sheer volume of detections it generates.

Indeed, a statistically significant—albeit extremely weak—correlation between time of detection and elevation was observed (Wald test, *p* = 1.8×10^−17^, linear regression *R*^2^ = 0.0065, slope = −5.59 s/m; Fig. 4c, *black line*). This corresponds to a delay in detections of roughly 16 minutes from the highest site (`kosci_north4; 2220` m) to the lowest site (`kosci_south0; 2046` m). We could tentatively take this as an indication that the bulk of the moths are moving downhill, although from this analysis alone, we are unable to disentangle that hypothesis from the hypothesis that moths at lower altitudes merely emerge from (and/or return to) their aestivation crevices later than higher-altitude moths (see Fig. 5 for illustration).

**Figure 5:**
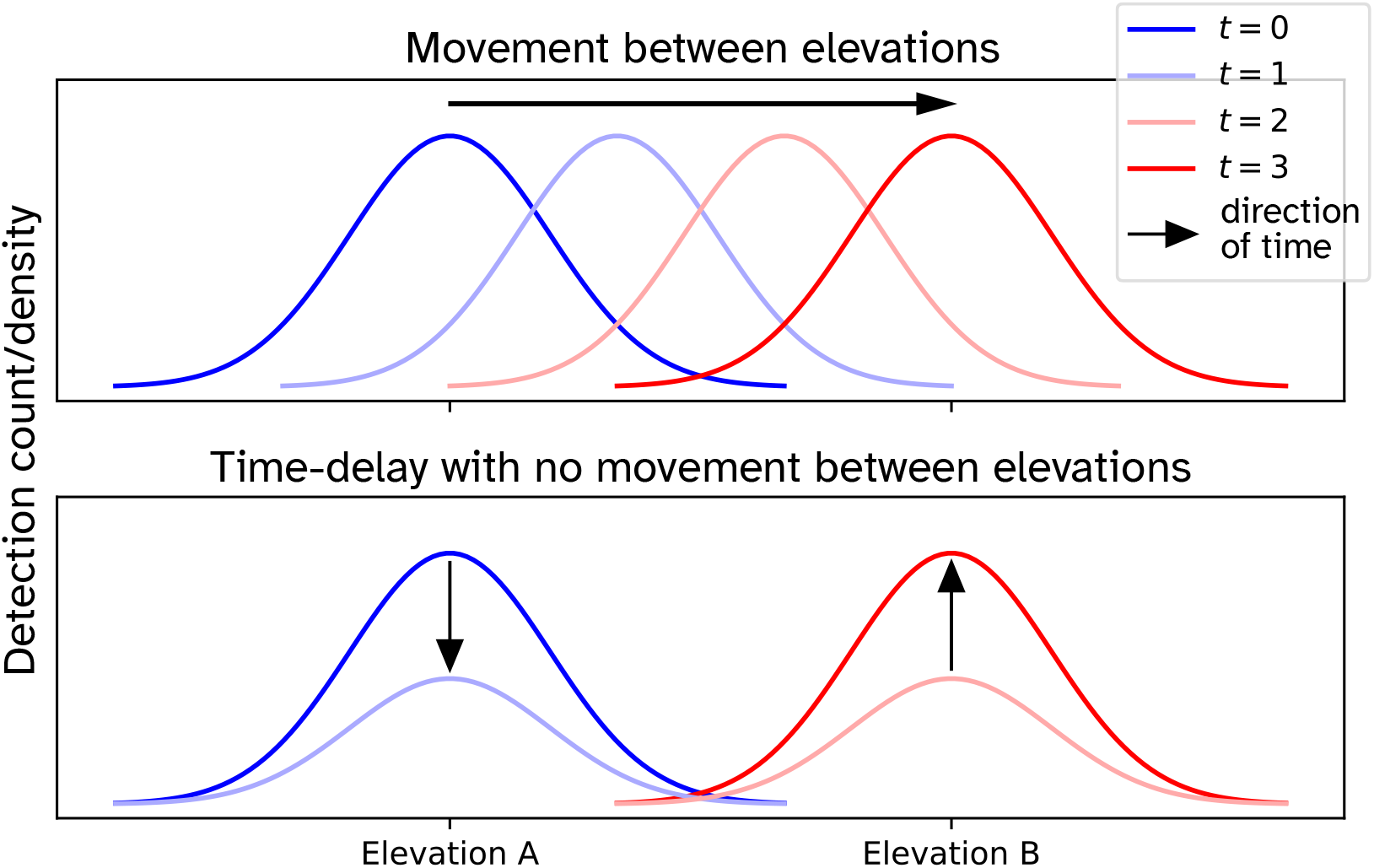
Scenarios which could lead to the observation that detection time depends on elevation. **Top panel:** In the first scenario, a lag in detection times is observed at elevation B, relative to elevation A, due to the movement of flying moths from elevation A to B. **Bottom panel:** In the second scenario, the lag is due to moths at elevation B emerging from—and returning to—their aestivation sites later than those at elevation A.

So far, we have attempted to detect the movement of Bogong moths along an altitudinal gradient by recording their location (i.e. displacement) over time. However, this analysis does not incorporate any information regarding the identity of the detected moths. This, along with the fact that moths may be present at a given location without being observable (for instance, a moth might not be airborne at a particular time), prevents us from concluding—with absolute certainty—that moths are indeed moving between elevations. If we knew that a particular moth had been detected at a given elevation, and detected again at another elevation a few minutes later, we could say with certainty that the moth moved between those elevations. Alas, there is barely enough information in the images taken by the wildlife cameras to positively identify species, let alone to identify individual moths that are members of a local population numbering in the millions.

### 3.2 Evidence of orientation behaviour

Displacement is, of course, not the only way to measure movement. We can also measure its derivative with respect to time; namely, velocity (the combination of direction of displacement, which we call “track direction,” and speed). In our case, we are only interested in the track direction of flights, which conveniently our method measures.

Flight track directions at each respective site in both transects showed significant departures from uniform circular distributions (Fig. 6), as determined by Moore’s modified Rayleigh tests (*p* < 0.05 for all locations, Moore, 1980). Furthermore, track directions from each pair of locations were signif-icantly different from each other, as determined by pairwise Mardia-Watson-Wheeler tests (*p* < 0.05 for each pair, Mardia, 1969). Thus, the flights of the Bogong moths were directed, and the direction of flight depended on location. This in itself is not surprising, however it is ethologically relevant, since directed movement requires behavioural control in response to external stimuli (Cheung et al., 2007).

**Figure 6:**
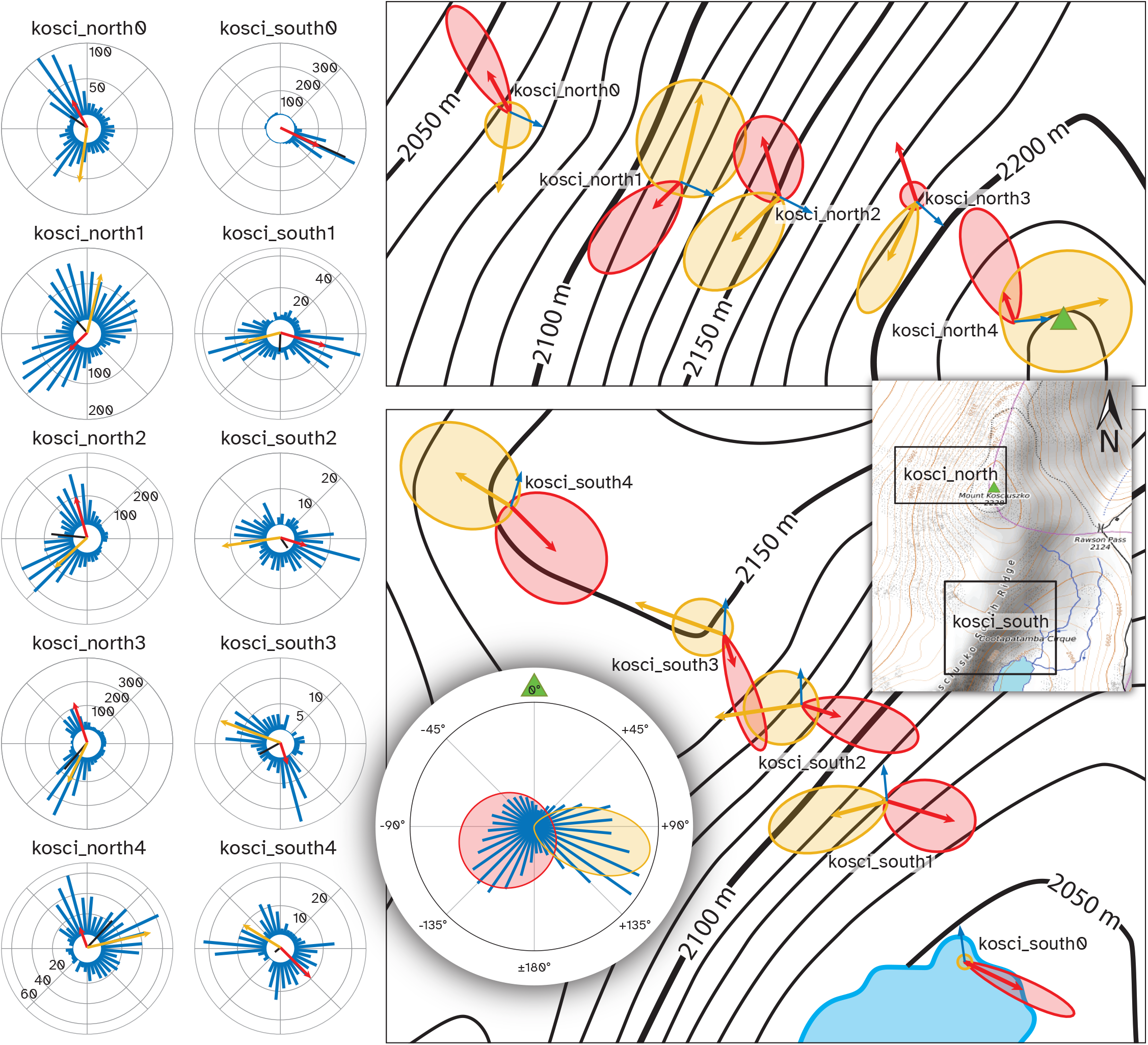
Distribution of flying insect track directions over the course of the evening of 18^th^ February 2021, by location. **Left panel:** *Blue bars* show histograms of track directions for the given location, with scale (counts) indicated on circular axes. *Black bars* show mean vector of all detections at the respective location. *Red arrows* show direction and weight of the first component of a bimodal von Mises distribution for the respective location, computed using the CircMLE R package (Fitak and Johnsen, 2017) and *yellow arrows* show the second component. Where only a *red arrow* is shown, the data were better explained by a unimodal distribution. **Top-right panel:** Arrows from the *left panel*, with von Mises probability density functions of each component also plotted. Plots are placed at their respective camera locations on a map of the `kosci_north` transect. *Blue arrows* show the azimuth of the summit of Mt. Kosciuszko from the respective locations. *Green triangle* shows summit of Mt. Kosciuszko. **Bottom-right panel:** Follows the conventions of the *top-right panel*, showing `kosci_south` transect. **Inset, right:** Map of Mt. Kosciuszko showing locations of top-right and bottom-right panels. *Green triangle* shows summit of Mt. Kosciuszko. **Circular inset, bottom:** Histogram (*Blue bars*) of flight track directions relative to the azimuth of the summit of Mt. Kosciuszko (*green triangle*) for all detections across all locations, shown with probability density functions of components of a bimodal von Mises model of the data (*red* and *yellow regions*).

Generally speaking, the distributions of flight track directions at each location were bimodal (Fig. 6; Table A.1) and the two modes were not separated by 180° (i.e. the bimodality of the directions was not a result of axially-directed flight). There was one notable exception to this trend, with moths detected at `kosci_south`0 showing a unimodal south-easterly flight track direction tendency (Fig. 6, southernmost site; Table A.1). Trends in flight track direction which were seen during nautical twilight (i.e. before 21:00 AEDT) were continued throughout the night, albeit with a much lower density of moths (Fig. A.2).

It is clear from our analyses that the Bogong moths exhibited orientation behaviour, although it is not immediately clear *how* they were orienting themselves. We know from laboratory assays of orientation behaviour that Bogong moths are able to orient themselves relative to visual landmarks in conjunction with the Earth’s magnetic field (Dreyer et al., 2018), and celestial cues (Dreyer and Adden et al., in prep.). Another possible source of directional information is wind, which is known to be used to control flight direction in another species of migratory noctuid moth, *Autographa gamma* (Chapman et al., 2008), and it is likely that Bogong moths also possess this ability. We wish to evaluate these possible sources of directional information with regard to our measurements of the orientation behaviour of Bogong moths in the wild.

Cues from celestial objects and the Earth’s magnetic field would be roughly the same across all study locations, given the small geographical area covered by the study (all locations were within 1.2 km of each other). Therefore, these cues alone could not explain the differences in the distributions of flight track directions observed across the locations (Fig. 6, *left panel*).

Wind speeds at 3 pm, 18^th^ February, and 9 am, 19^th^ February 2021 were moderate to fresh—4 ms^-1^ easterly and 5 ms^-1^ north-northwesterly, respectively (recorded at Thredbo Top Station, circa 4.6 km from the summit of Mt. Kosciuszko; Bureau of Meteorology, 2021). These wind speeds are similar to the likely airspeed of a motivated Bogong moth, and could have an important impact on their resultant track direction, especially for high-flying moths. The speed and direction of wind can be modulated by topography, so it is possible that wind varied between the study locations, although this was not measured, so we cannot rule out the possibility that wind could explain the observed differences in flight track directions across the locations. However, bimodal distributions of flight track directions would be hard to explain with wind alone.

We can say with certainty that the terrestrial visual panorama varies greatly across the study locations, especially between the two transects, which are on opposing sides of the highest point in the mountain range. Dreyer et al. (2018) showed Bogong moths orienting themselves relative to the azimuth of an abstraction of the silhouette of a mountain peak (namely, a black triangle on a white background, above a black horizon). Mountains are striking visual landmarks in the Australian Alps (Paterson, 1890)^3^ where Bogong moths spend their summer. It is therefore reasonable to predict that wild Bogong moths exhibiting directed flights in their summer range would fly in directions relative to mountain peaks. As it happens, the summit of Mt. Kosciuszko (2228 m) is the highest peak in Australia, and it is also the closest peak to all of the study locations in both transects. We therefore proceed by comparing the azimuth of the summit of Mt. Kosciuszko with flight track directions of the Bogong moths we detected.

Indeed, flight track directions relative to the azimuth of the summit of Mt. Kosciuszko clustered bimodally (pooled across all locations, confirmed by AIC-based maximum-likelihood model selection, Table A.3; distributions shown in Fig. 6, *right panels* and *circular inset*). The respective means of the two components of a bimodal von Mises model, fit to flight track directions relative to the azimuth of the summit were −118° (SD: 80.2°) and +103° (SD: 28.8°) (Fig. 6, *circular inset*). As both of these are greater (in absolute terms) than 90°, the Bogong moths were, in aggregate, moving away from the summit. And since there is no higher point in Australia than the summit of Mt. Kosciuszko, the moths were also moving downhill, on paths which would take them around the mountain.

### 3.3 Why do aestivating Bogong moths take flight?

Our analyses of the rich dataset produced by our new method have so far told us *what* the Bogong moths were doing (moving downhill), and have provided us with a robust hypothesis for *how* they were doing it (flying relative to the azimuth of the nearest—and highest—summit). What remains to be answered is, *why* do Bogong moths behave this way? Indeed, why do they take flight almost every evening throughout summer—a decidedly non-dormant activity—when they are supposedly aestivating?

We have presented evidence from both displacement and velocity data indicating that on the evening of 18^th^ February 2021, Bogong moths on Mt. Kosciuszko were, in aggregate, moving downhill. This movement was characterised not by a straight-line departure from the peak of the mountain, nor a departure in a particular direction. Instead, it was characterised by motion *relative* to the azimuth of the peak, with moths presumably fixing the direction of their flight by holding the azimuth of the summit at a constant obtuse angle, with respect to their direction of travel, leading to trajectories that would resemble portions of outward logistic spirals centred on the summit, when viewed from above.

Interestingly, this *almost* matches qualitative observations made by JRAW and EJW from the same vantage point near `kosci_north`2, about 14 months earlier. An excerpt from JRAW’s field notes from 20:45 on the 28^th^ December 2019 reads,

“When I look up the hill, I can see fast-moving moths moving right to left [and left to right]. And when I look to the side, along the mountain, the overarching movement is a slow movement uphill. There seems to be two different modes of moth flight—there’s a slow upward movement, and then a fast lateral movement in both directions.”

Perhaps we were seeing the equivalent pattern of flight directions to those on 18^th^ February, 2021, with an uphill rather than downhill trend (in this case, flight directions would form an acute angle with the azimuth of the summit, resulting in inward-logistic-spiral trajectories).

If this is true, an appealing explanation for the up-and downhill movements is that Bogong moths were seeking new aestivation sites of higher elevation on 28^th^ December 2019, while on 18^th^ February 2021, the moths on Mt. Kosciuszko were getting ready to leave. This would make sense, as temperatures typically don’t peak until January in Australia, so it is likely that Bogong moths are still in the forward half of their round-trip migration in late December (and higher elevations have lower temperatures, thanks to adiabatic expansion). Notably, daytime temperatures in the Australian Alps were high in the last few days of 2019, reaching 23.3°C on 28^th^ December at Thredbo Top Station (Bureau of Meteorology, 2019),^4^ while on 18^th^ February 2021, the temperature only reached a more moderate 15.6°C at the same location (Bureau of Meteorology, 2021). We know from long-term monitoring data that the bulk of the Bogong moths had already left the lower elevation aestivation sites of Mt. Gingera and Ken Green Bogong by mid-February 2021, and that numbers on Mt. Kosciuszko were declining in that month (Wallace et al., 2022), so it is reasonable to conclude that the return migration had begun.

A possible explanation for why Bogong moths move laterally (i.e. in a logistic spiral) around the mountain, rather than in a straight line, is that in addition to altering elevation, these flights are used to calibrate the moths’ internal compasses. The flights only occur just after sunset, so the flying moths would be able to see the azimuth of the sunset, which is an extremely stable compass cue. Similarly, they could be using their magnetic sense to perceive the Earth’s magnetic field (Dreyer et al., 2018), an even more stable compass cue. Meanwhile, the moths could be taking snapshots of the terrestrial, and possibly celestial panorama (as suggested for dung beetles, el Jundi et al., 2016), which they could later use as a terrestrial compass cue (Zeil, 2012) while they remain in the area, helping them to navigate at the start of their return migration to their breeding grounds. There is a distinct possibility that each of these cues are taken together, and these evening flights are used by Bogong moths to calibrate multi-sensory internal compasses which they eventually use for their return migration. Such multi-sensory compass calibrations are thought to be performed by migratory songbirds (reviewed by Foster et al., 2018; Pakhomov and Chernetsov, 2020).

In order to see the entire distant terrestrial panorama in the direction of their return migration, a moth would either have to fly up above the summit of the mountain, or it would have to traverse around the summit while taking snapshots, since approximately half of the panorama would be occluded by the mountain for a moth flying below the summit. The latter would likely be a safer strategy, as a higher altitude flight could present a risk of strong winds blowing the moth away from the mountain entirely, forcing the moth to expend more energy to return if it is not yet ready to leave. This process would also be useful for the forward phase of Bogong moth migration, as it would enable the moths to check the horizon for taller, and therefore more favourable (particularly in late summer) mountains, which they could then orient relative to in the subsequent leg of their journey, akin to beacon-aiming performed by wood ants (Graham et al., 2003).

## 4 Additional remarks

Some of the most interesting behavioural phenomena involve highly complex and fragile mechanisms which are easily disturbed, rendering them challenging to study. Animal migration is a conspicuous example of such fragility. For instance, merely eclosing otherwise wild Monarch butterflies in captivity is enough to disrupt their migratory orientation behaviour (Tenger-Trolander et al., 2019). Therefore, it is important that we are able to support laboratory results of animal behaviour with data obtained in the wild, ideally without any potentially disruptive manipulations (i.e. exposing the animal only to natural stimuli). In this paper, we have shown that for certain questions, our new method— which enables us to inexpensively make ethological observations of wild insects in a high-throughput, quantitative, and completely non-invasive manner—allows us to do just that. Conversely, laboratory-based experimentation is extremely useful for testing specific hypotheses, as it allows us to present animals with controlled stimuli of our own choosing (we could, for instance, change the azimuth of a prominent landmark).

Our results have generated a number of interesting and plausible hypotheses relating to the previously unexplained summer evening flight behaviour of Bogong moths. The first hypothesis is that the evening flights serve a specific navigational purpose. In particular, these flights might be used by Bogong moths to calibrate their internal compasses by integrating directional information provided by the azimuth of the setting sun, the geomagnetic field, and the visual panorama. Second, the visual panorama might be used by the moths on both the forward and return legs of their migration.

On the forward leg, it would enable them to identify other mountains which may be more suitable for continuing their aestivation on (e.g. higher mountains). On the reverse leg, it could be used as a reliable compass, which provides valid directional information for moths remaining in the local area. Such a compass would be especially useful for helping the moths select favourably-directed winds for their return migration (Chapman et al., 2008). Third, to navigate effectively, the Bogong moths may need to see the visual panorama in the direction of their migration, and to access this, they fly around the nearest prominent mountain peak, rather than flying to a high altitude, where there is a risk of being blown off course.

Mouritsen (2018) listed twenty of the most important open questions in long-distance navigation research for the next twenty years. Included in this list is the question,

“How does the pinpointing-the-goal phase work in a Monarch butterfly or Bogong moth, which can pinpoint their very specific wintering [or, for the Bogong moth, summering] locations even though they have never been there before?”

If our hypothesis that Bogong moths use their summer evening flights during their forward migration to identify taller and taller mountains turns out to be true, then this could go some way to answering this question. The Bogong moths may have never had an opportunity to learn where the tallest mountains in the Australian Alps are, but by employing a beacon-aiming navigational strategy they could simply find out when they get there. Of course, there must be other factors at play as well, since Bogong moths don’t all end up on the highest peak (i.e. Mt. Kosciuszko). Lower peaks with otherwise favourable conditions (e.g. with availability of crevices with suitable temperature, humidity, and darkness) could also end the forward migration.

## 5 Author contributions

EJW, JZ, and JRAW conceived the project. EJW and JRAW designed the experiment. JRAW collected the data, devised the algorithms, and wrote the software. DD and JRAW analysed the data. JRAW wrote the first draft of the manuscript. All authors critically interpreted the results, and contributed to the writing and editing of the manuscript.

## 6 Acknowledgements

EJW and JRAW are grateful for funding from the European Research Council (Advanced Grant No. 741298 to EJW), and the Royal Physiographic Society of Lund (to JRAW). JRAW is thankful for the support of an Australian Government Research Training Program (RTP) Scholarship. EJW holds Scientific Permits for collection and experimental manipulations of Bogong moths in several alpine national parks and nature reserves (NSW Permit SL100806). We are extremely grateful to Mark Rullo for assisting with the fieldwork, and to Dr. Stanley Heinze for useful discussions and providing computational resources.

## A Appendix

**Table A.1:** Circular distribution models and corresponding output parameters selected using Akaike’s information criterion (AIC), computed using the CircMLE R package (Fitak and Johnsen, 2017) on flying insect detections at the respective camera locations (Loc.; for brevity, “kosci_” prefixes are removed from each location name). Model selection was performed on the models defined by Schnute and Groot (1992). Models appearing in table: 2B = “symmetric modified unimodal,” 5A = “homogeneous bimodal,” 5B = “bimodal.” Models are mixtures of von Mises distributions with two components *i* (*i* = 1, 2). φ_*i*_ denotes the mean direction of component *i* (in radians), κ_*i*_ the von Mises concentration parameter of component *i*, and λ the proportion assigned to the first component. θ is the azimuth of the summit of Mt. Kosciuszko (the nearest and highest peak) from the respective location. _†_Parameter fixed by model (λ = 0.5, κ_2_ = 0). _‡_Concentration parameters are assumed equal by model (κ_1_ = κ_2_).

| Loc. | Model | $\varphi_1$ | $\kappa_1$ | $\lambda$ | $\varphi_2$ | $\kappa_2$ | $\theta$ |
| --- | --- | --- | --- | --- | --- | --- | --- |
| north0 | 5B | 5.782 | 16.056 | 0.365 | 3.286 | 0.584 | 1.993 |
| north1 | 5B | 3.941 | 6.111 | 0.302 | 0.234 | 1.103 | 1.972 |
| north2 | 5B | 6.001 | 2.079 | 0.490 | 3.965 | 3.589 | 2.037 |
| north3 | 5B | 5.960 | 0.628 | 0.487 | 3.592 | 13.697 | 2.303 |
| north4 | 5B | 5.929 | 8.383 | 0.250 | 1.342 | 1.166 | 1.493 |
| south0 | 2B | 2.019 | 46.502 | 0.5<sup>†</sup> | - | 0<sup>†</sup> | 6.148 |
| south1 | 5B | 1.859 | 2.235 | 0.547 | 4.457 | 5.203 | 6.189 |
| south2 | 5B | 1.870 | 9.389 | 0.320 | 4.567 | 0.642 | 6.255 |
| south3 | 5B | 2.826 | 31.053 | 0.265 | 5.060 | 1.037 | 0.045 |
| south4 | 5A | 2.356 | 1.920<sup>‡</sup> | 0.495 | 5.263 | 1.920<sup>‡</sup> | 0.292 |

### A.1 Model selection tables

**Table A.2:**
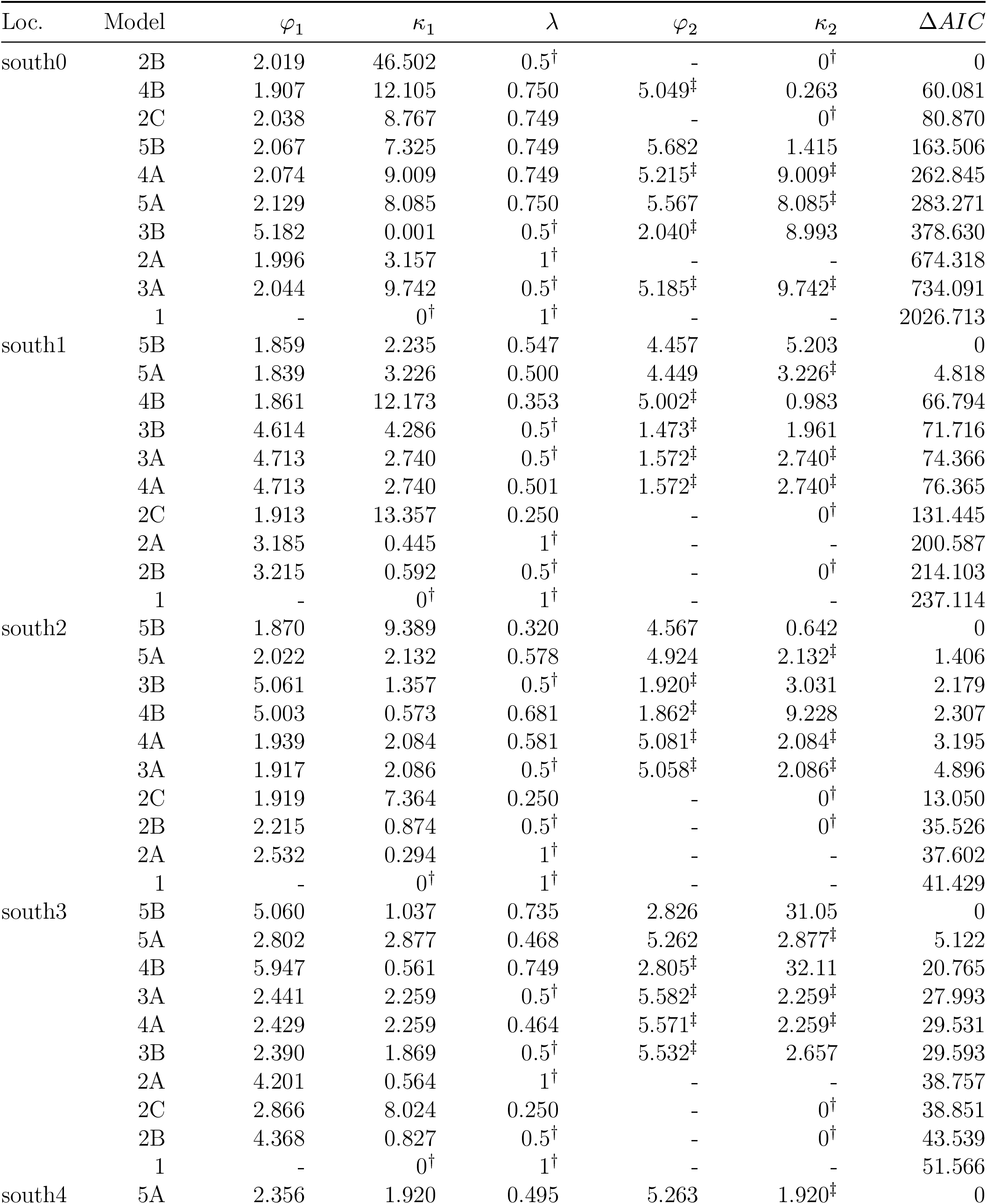
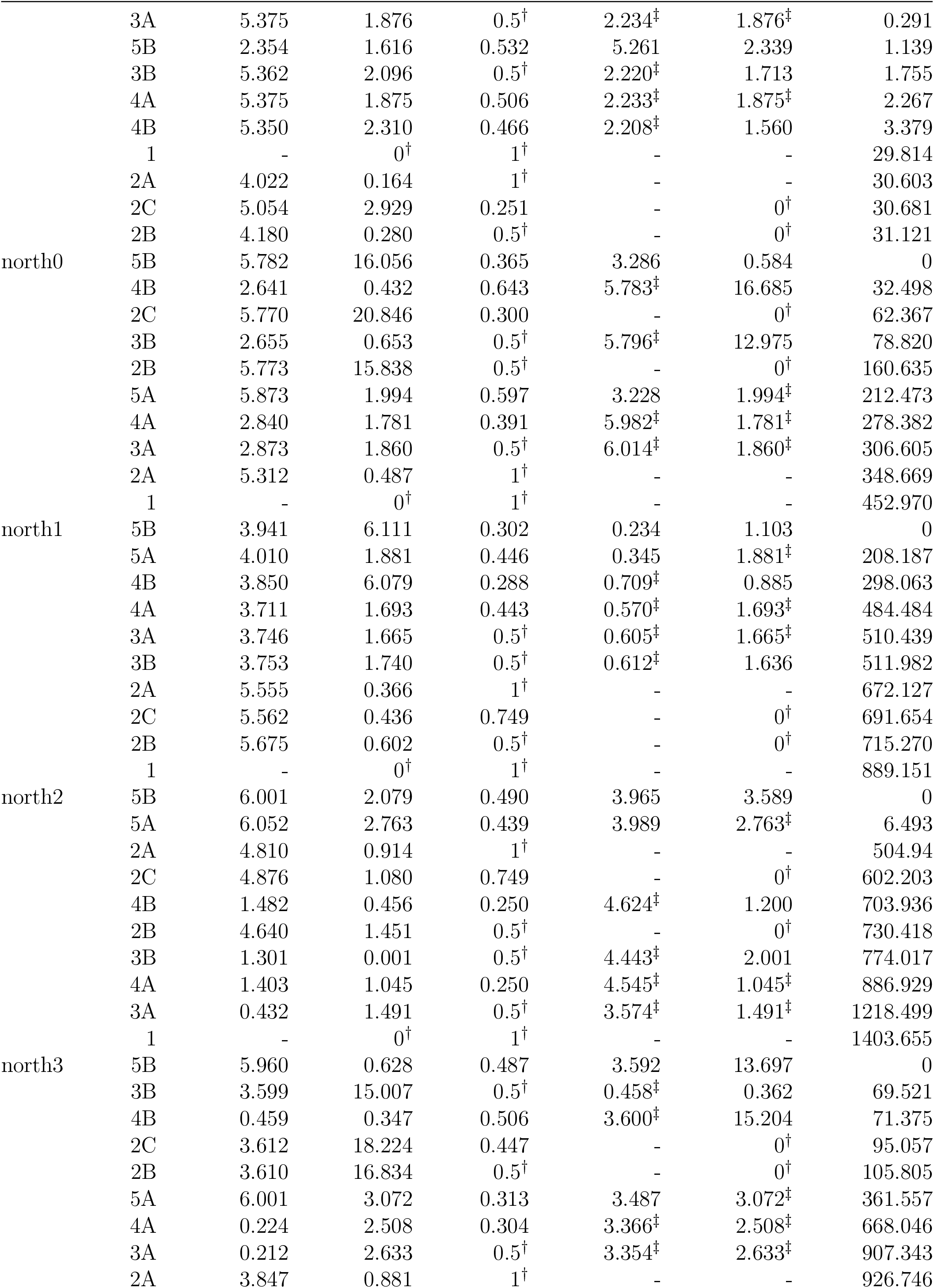
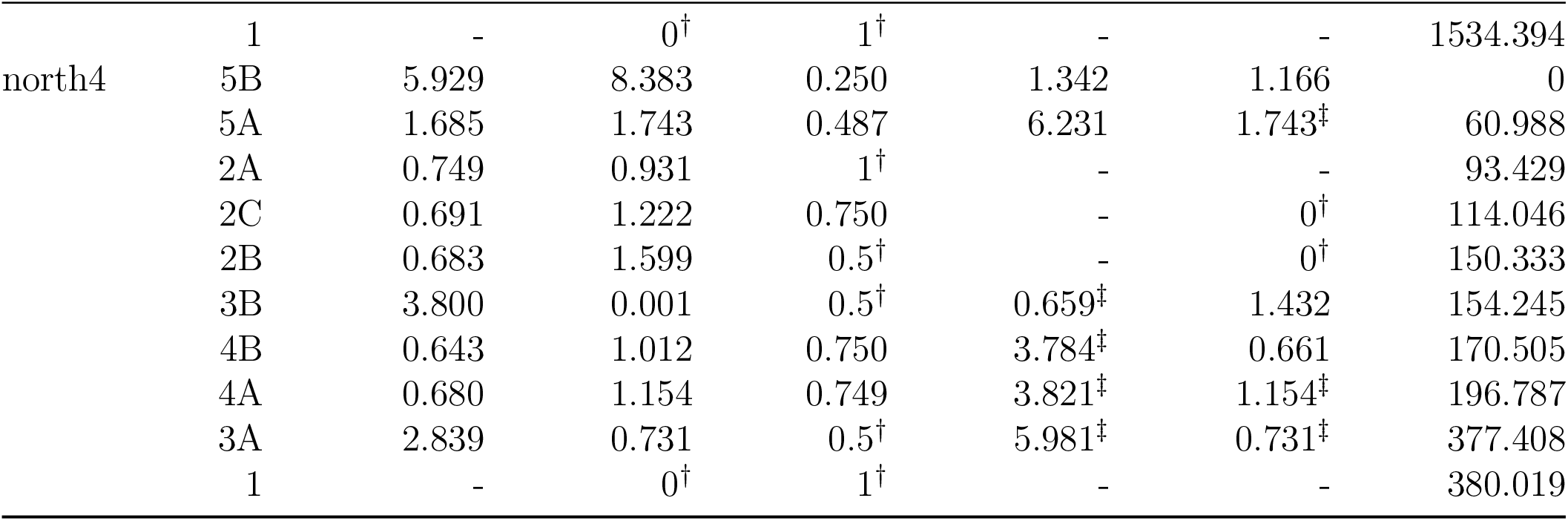
Model selection table for track directions at each camera location. ^†^Parameter fixed by model. ^‡^Parameter depends on another parameter in model (i.e. φ_2_ = φ_1_ + π(mod 2π), or κ_1_ = κ_2_). Models for each location are sorted by the model selection criterion, Δ*AIC*. All other parameters follow the conventions of Table A.1.

**Table A.3:** Model selection table for track directions relative to the azimuth of the summit of Mt. Kosciuszko. Follows conventions of Table A.2.

| Model | $\varphi_1$ | $\kappa_1$ | $\lambda$ | $\varphi_2$ | $\kappa_2$ | $\Delta AIC$ |
| --- | --- | --- | --- | --- | --- | --- |
| 5B | 2.055 | 0.741 | 0.654 | 4.480 | 4.584 | 0 |
| 5A | 4.463 | 1.650 | 0.561 | 1.899 | $1.650^\ddagger$ | 385.854 |
| 4B | 4.466 | 5.483 | 0.301 | $1.325^\ddagger$ | 0.393 | 693.199 |
| 2C | 4.369 | 7.320 | 0.251 | - | $0^\dagger$ | 897.322 |
| 3B | 4.549 | 2.412 | $0.5^\dagger$ | $1.408^\ddagger$ | 0.987 | 1012.238 |
| 4A | 4.607 | 1.429 | 0.582 | $1.465^\ddagger$ | 1.429 | 1238.613 |
| 2A | 3.645 | 0.410 | $1^\dagger$ | - | - | 1368.769 |
| 3A | 1.537 | 1.483 | $0.5^\dagger$ | 4.679 | 1.483 | 1387.302 |
| 2B | 3.851 | 0.802 | $0.5^\dagger$ | - | $0^\dagger$ | 1442.646 |
| 1 | - | $0^\dagger$ | $1^\dagger$ | - | - | 2274.647 |

### A.2 Luminance recordings

**Table A.4:** Luminance recordings from various locations inside and outside a Bogong moth aestivation cave, in February, 2021. Recordings were taken using a digital photometer (Hagner, model ERP-105).

| Location | Time | Reading | Multiplier |
| --- | --- | --- | --- |
| Inside cave entrance | 20:22 18/02 | 100 | 0.01 |
| (recorded off standard) | 20:35 18/02 | 78 | 0.01 |
|  | 20:56 18/02 | 14 | 0.01 |
| Outside cave, | 20:22 18/02 | 442 | 10 |
| (recorded off standard) | 20:35 18/02 | 250 | 1 |
|  | 20:56 18/02 | 490 | 0.01 |
| Main front wall inside cave | 20:30 18/02 | 17 | 0.01 |
| (no moths) | 20:33 18/02 | 41 | 0.01 |
|  | 06:51 19/02 | 461 | 0.01 |
| Cave, far right | 20:30 18/02 | 6 | 0.01 |
| (opposite moths) | 06:51 19/02 | 22 | 0.01 |
| Cave, right, left of rock | 20:31 18/02 | 2 | 0.01 |
| (opposite moths) | 06:52 19/02 | 14 | 0.01 |
| Cave, back, shaded | 20:32 18/02 | 1 | 0.01 |
| (no moths) | 06:52 19/02 | 34 | 0.01 |
| Cave, far left (no moths) | 20:32 18/02 | 12 | 0.01 |
| Cave, left (opposite moths) | 06:53 19/02 | 9 | 0.01 |

**Figure A.1:**
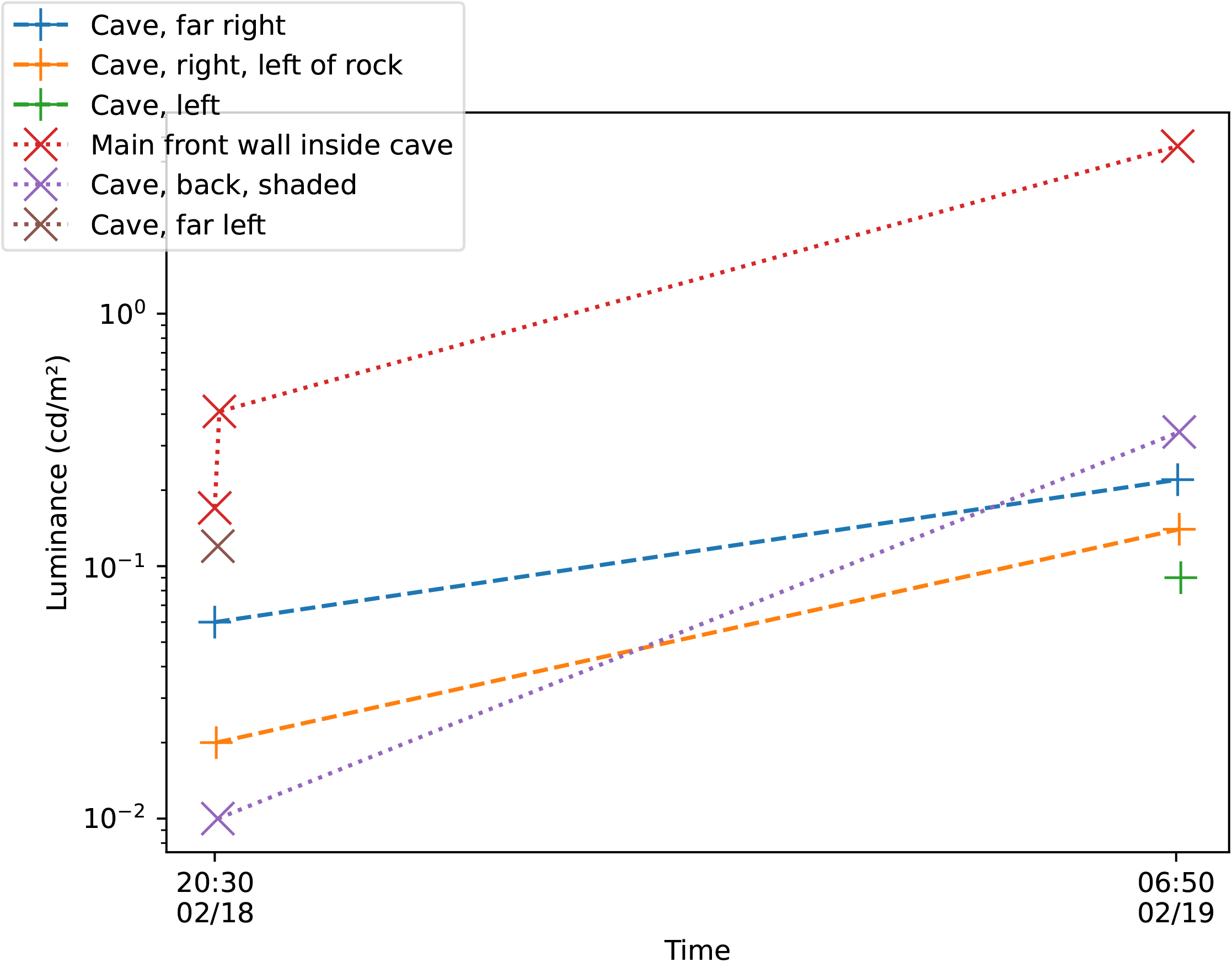
Occupied positions within a Bogong moth aestivation cave on Mt. Kosciuszko (“+” markers, connected with *dashed lines*) tend to be darker than unoccupied positions (“×” markers, connected with *dotted lines*), particularly during the day (right side of plot), as measured by digital photometer (Hagner, model ERP-105).

### A.3 Flight track orientations

**Figure A.2:**
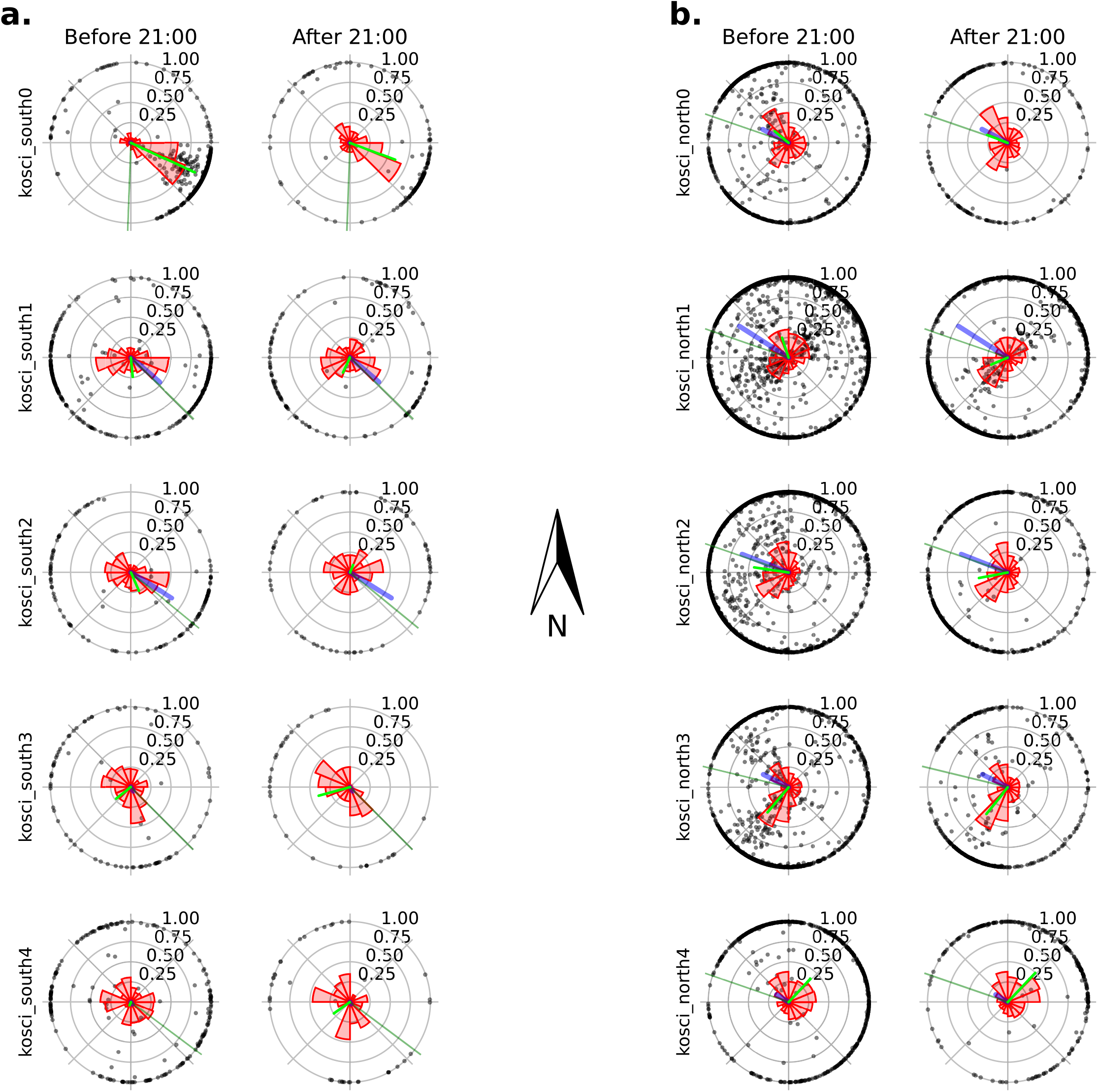
Trajectories of detected insects during nautical twilight (before 21:00 AEDT) and after for both transects. Columns indicate time period and rows indicate location. **a**. `kosci_south` transect. **b**. `kosci_north` transect. *Black dots*: Track (direction of displacement) of detected insect trajectories. Radius indicates the straightness of the trajectory, calculated as distance travelled divided by displacement (in pixel units). *Red bars*: Circular histogram of detected insect trajectories. The bars are equiareal (area—not height—indicates proportion of detections contained within each bin). *Lime green line*: Mean track (direction of displacement) of detected insect trajectories. Radius indicates circular mean vector length (with values closer to one indicating more concentrated tracks). *Blue line*: Fall line of the slope at the position of the camera. The direction indicates the direction of maximum gradient (perpendicular to topographic lines), and the radius indicates the gradient itself. *Dark green line*: Indicates the bearing of the base of the camera.

## Footnotes

1 To illustrate the point, a note from JRAW’s field book recounting one of these evening flight events reads, “I decided to see if it was enough to simply reach out an open hand and close it into a fist in order to catch a moth. It worked—first try.”

2 This information does not extend to the direction the insect is flying with respect to the camera, but it does include the orientation of flight (with 180° ambiguity).

3 From *The man from Snowy River* (Paterson, 1890), “And down by Kosciusko, where the pine-clad ridges raise Their torn and rugged battlements on high, …”

4 and 24.6°C on the 30^th^, and then 26.5°C just four days later (4^th^ January 2020) which was the worst day for Kosciuszko National Park (KNP) of the 2019-2020 bush fire season, which saw over 200,000 ha of KNP burn.

